# Revisiting typing systems for group B *Streptococcus* (GBS) prophages: an application in prophage detection and classification in GBS isolates from Argentina

**DOI:** 10.1101/2024.05.08.593127

**Authors:** Verónica Kovacec, Sabrina Di Gregorio, Mario Pajón, Chiara Crestani, Tomás Poklepovich, Josefina Campos, Uzma Basit Khan, Stephen D. Bentley, Dorota Jamrozy, Marta Mollerach, Laura Bonofiglio

## Abstract

Group B *Streptococcus* (GBS) causes severe infections in neonates and adults with comorbidities. Prophages have been reported to contribute to GBS evolution and pathogenicity. However, no studies are available to date on the presence and diversity of prophages in GBS isolates from humans in South America. This study provides insights into the prophage content of 365 GBS isolates collected from clinical samples in the context of an Argentinean multicentric study. Using whole genome sequence data, we implemented two previously proposed methods for prophage typing: a PCR approach (carried out *in silico*) coupled with a blastx-based method to classify prophages based on their prophage group and integrase type, respectively. We manually searched the genomes and identified 325 prophages. However, only 80% of prophages could be accurately categorised with the previous approaches. Integration of phylogenetic analysis, prophage group and integrase type allowed for all to be classified into 19 prophage types, which correlated with GBS clonal complex grouping. The revised prophage typing approach was additionally improved by using a blastn search after enriching the database with 10 new genes for prophage group classification combined with the existing integrase typing method. This modified and integrated typing system was applied to the analysis of 615 GBS genomes (365 GBS from Argentina and 250 from public databases), which revealed 29 prophage types, including 2 novel integrase subtypes. Their characterization and comparative analysis revealed major differences in the lysogeny and replication modules. Genes related to bacterial fitness, virulence or adaptation to stressful environments were detected in all prophage types. Considering prophage prevalence, distribution and their association with bacterial virulence, it is important to study their role in GBS epidemiology. In this context, we propose the use of an improved and integrated prophage typing system suitable for rapid phage detection and classification with little computational processing.

**Author summary:** Bacteriophages, which are viruses that infect bacteria, exert a profound influence on microbial evolution when integrated into the bacterial genome, a state in which they are called prophages. It has been proposed that prophage acquisition played a role in the emergence of Streptococcus agalactiae (GBS) as a human pathogen in European countries. Further study and characterization of prophages of GBS from around the world would provide valuable insights into the mechanisms underlying GBS adaptation, evolution and epidemiology.

Unfortunately, existing tools for prophage screening exhibit limitations in the detection and classification of all prophages present in GBS genomes. To address this issue, in this work we propose a new prophage typing system that allows the detection and classification of GBS prophages based on both their phylogenetic lineage and integration site within the bacterial genome. Using this methodology we were able to identify 29 prophage types in 615 GBS isolated globally. We further characterised these prophages and found that they carried genes that could give an evolutionary advantage to their host and that different lineages of GBS carried different prophage types.

Comprehensive exploration and characterization of prophages represent an indispensable endeavour, providing critical insights into microbial evolution, epidemiology, and potential therapeutic interventions.

## Introduction

*Streptococcus agalactiae* (group B *Streptococcus*; GBS) is a commensal bacterium that colonises the human intestinal and genitourinary tracts. GBS is a major cause of neonatal sepsis and other perinatal infections, such as meningitis and pneumonia, globally [1]]. In recent decades, invasive infections caused by GBS in non-pregnant adults have been on the rise, especially in the elderly people and in those suffering from underlying medical conditions [[2–4]].

Prophages are important vehicles for horizontal gene transfer (HGT) [[5] and can constitute up to 20% of a bacterial genome. Prophages play a significant role in bacterial evolution by introducing genes that enhance bacterial fitness and virulence [6,7]. Furthermore, pathogenic strains tend to carry more phage-related genes than non-pathogenic strains [8–10], which was also observed for GBS [11].

GBS temperate bacteriophages (lysogenic prophages) were first described in 1969 in strains from bovine origin [12]. Recent studies on human GBS isolates revealed an association between certain prophages (some of possible animal origin) and the emergence of specific pathogenic GBS clones among isolates recovered from neonates and adults in Europe [11,13–15]. Little is known about the epidemiology of GBS prophages and their impact on pathogenicity in other geographical areas. To date, there are no reports of prophages in GBS isolates from South America [16].

Two approaches have been previously developed for the screening and classification of prophages in GBS genomes, one based on full-prophage sequence diversity [11] and the other based on integrase typing [16]. However, both have limitations and may under or overestimate prophage presence.

This study aims to analyse the prophage content in GBS genomes from Argentina by integrating and improving the existing methods for the detection and typing of GBS prophages, providing a novel strategy for a global surveillance of GBS prophage epidemiology and diversity.

## Materials and Methods

### M1. Isolate collection

We collected 450 GBS isolates from maternal carriage as well as invasive and urinary tract infections as part of a national multicentric study that involved 40 health centres in 12 provinces of Argentina between 2014-2015. All isolates had been characterised phenotypically (antibiotic susceptibility and serotyping) and invasive isolates had also been characterised genotypically (PFGE) [4,17,18].

### M2. Whole genome sequencing and data processing

Genomic DNA of 450 GBS isolates was extracted using a QIACube HT protocol and sequenced at the Wellcome Sanger Institute on Illumina NovaSeq 6000 platform (as part of our collaboration with the Juno consortium, https://www.gbsgen.net/). For 10 GBS isolates, genomic DNA was extracted using Wizard® Genomic DNA Purification Kit (Promega) and sequenced at the Malbrán Institute on Illumina MiSeq. Quality of the reads was assessed with FastQC v0.11.7 [19] and Kraken v0.10.6 [20]. *De novo* assemblies were obtained with SPAdes v3.12.0 [21] and quality checked with Quast v5.0.0 [22]. The 365/450 assemblies that passed the quality controls were annotated with Prokka v1.12 [23]. MLSTs were determined with the software mlst v2.22.1 [24] and assigned to clonal complexes [CC] using the PubMLST website [25,26] (https://pubmlst.org/organisms/streptococcus-agalactiae).

### M3. Prophage detection and typing

The methodology followed for the detection and typing of the GBS prophages is summarised in this section and Fig 1. For detailed information see M3 in S1 File.

**Fig 1.**
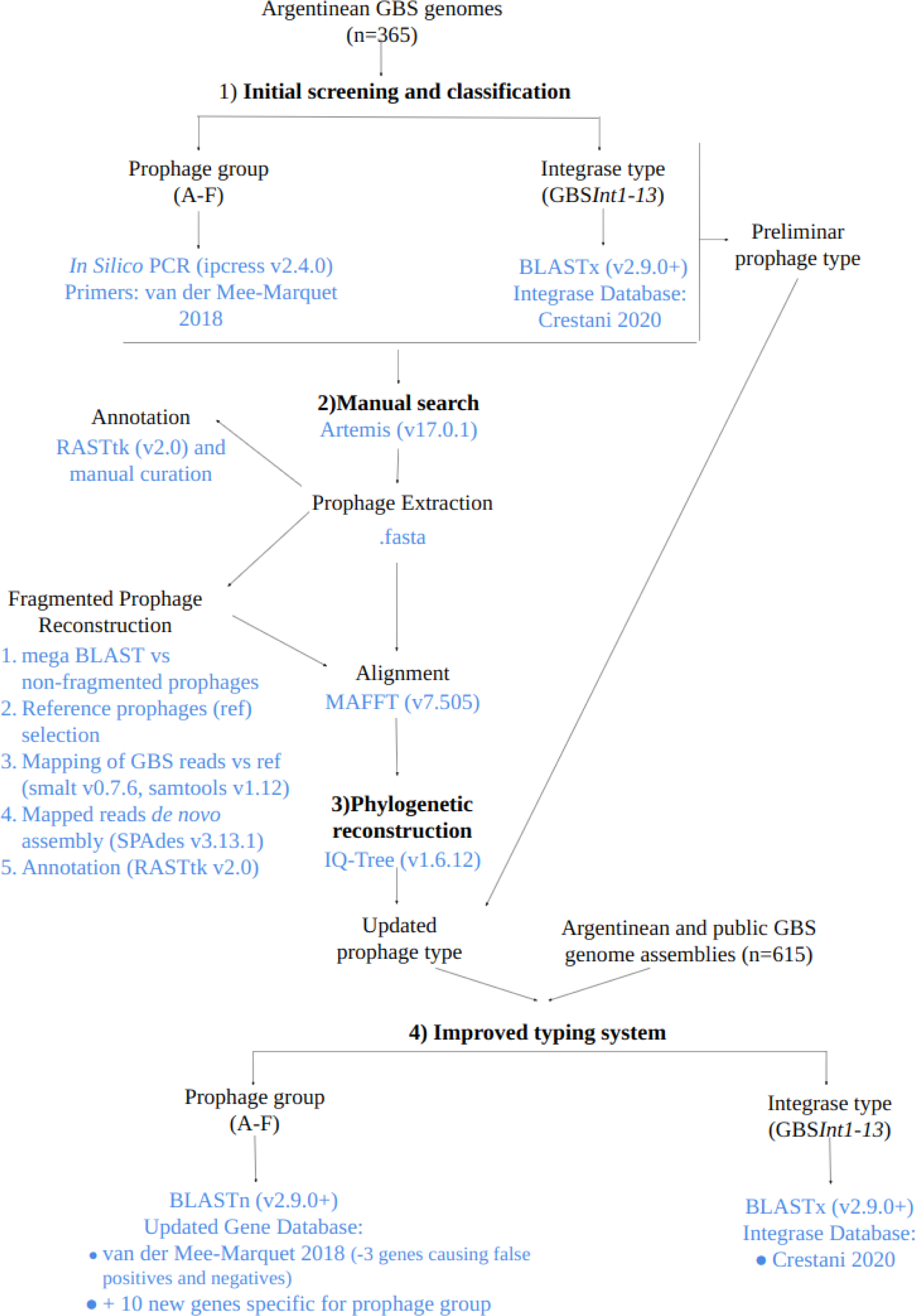
Summarised methodology used for GBS-prophage detection, typing and improvement of the prophage typing system.

In the first instance, prophage sequences were detected and classified using the previous screening methods for prophage groups [11] (here perfomed *in silico*) and integrase type [16]. Results of the two methods were combined and the putative prophages were preliminarily classified into prophage types according to prophage group and integrase type (Fig 1).

Prophage sequences within the assembled genomes were manually searched and extracted. Prophages fragmented across contigs were reconstructed by *de novo* assembly against reference prophages. All prophage sequences were annotated. (Fig 1).

The extracted prophage sequences were aligned and a phylogenetic tree was reconstructed. Prophages that could not be assigned to a prophage group during the initial screening stage were classified with the same group as the prophages in their phylogenetic cluster. Classification by prophage type was updated accordingly (Fig 1).

### M4. Improvement of the prophage typing system

The methods followed for the improvement of the prophage typing system are summarised in this section and Fig 1. For detailed information see M4 in S1 File.

In order to avoid false positive and false negative results obtained in the initial screening, we improved the detection of prophage groups by performing a blastn search against a curated database of prophage group-specific genes (Table S1 in S2 File). The methodology was tested on the 22 prophages detected by van der Mee-Marquet et al. (2018), all prophages from Argentinean GBS genomes and 615 GBS complete genomes from Argentina and public databases (Table S2 in S2 File). A result was considered positive when at least one of the genes for the prophage group was detected with a minimum of 75% of identity and coverage. Prophage types were then defined combining these results with those of integrase types.

### M5. Prophage characterization

The steps followed for prophage characterization are summarised in this section and Fig 2. For detailed information see M5 in S1 File and results section R2.

**Fig 2.**
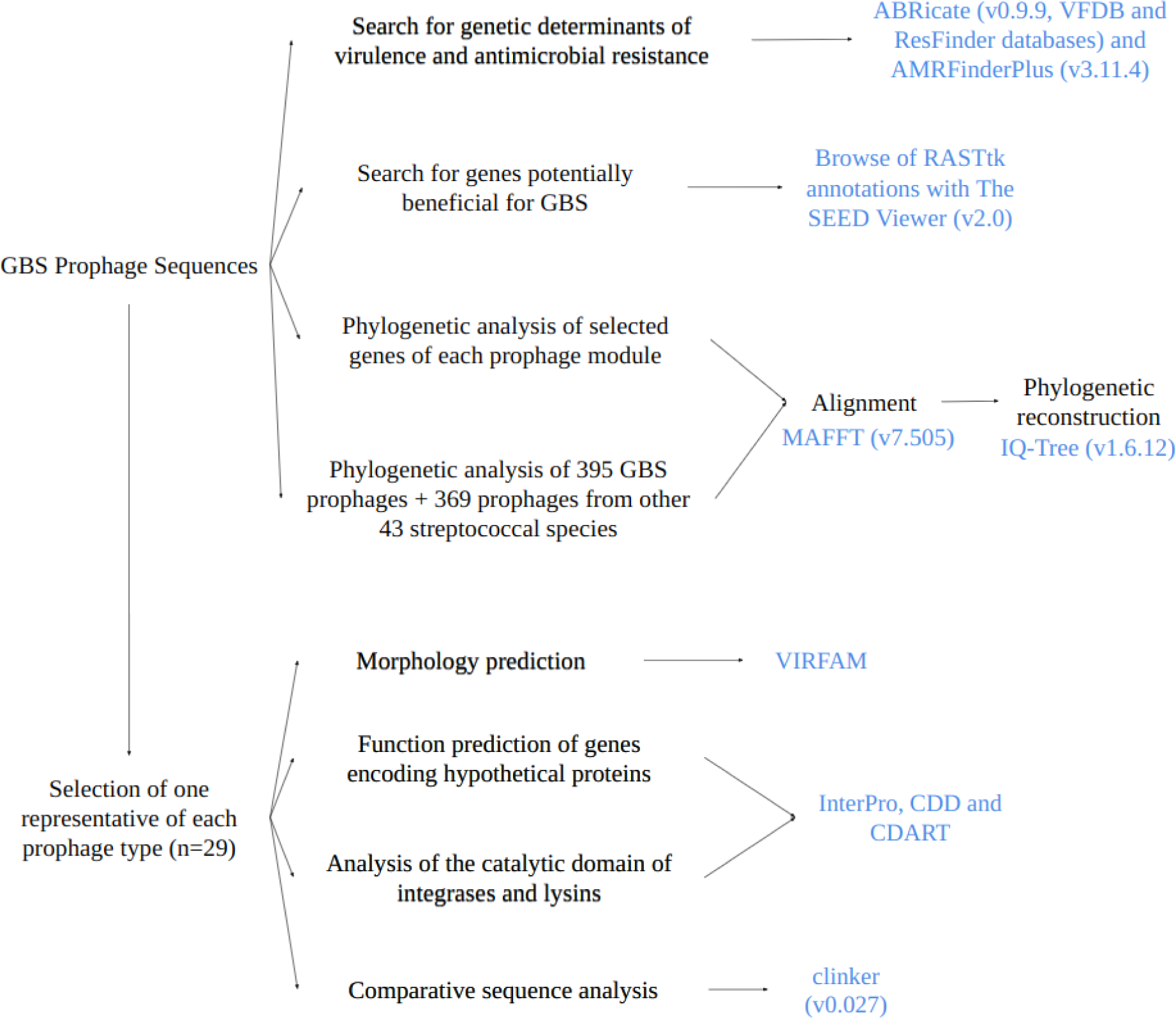
Summarised methodology used for GBS-prophage characterization.

Prophage sequences were searched for genetic determinants of virulence and antimicrobial resistance, as well as any genes potentially beneficial for the host bacteria. Genes coding for integrase, helicase, terminase large subunit, major capsid protein and lysin, were used for the phylogenetic analysis of each prophage module.

One phage of each prophage type (n=29, see results section) was selected for further characterization. The morphology of the prophages was predicted, as well as the function of the genes annotated as encoding hypothetical proteins and the catalytic domain of the putative integrases and lysins present. A comparative sequence analysis was performed to study the genetic differences between prophages of the same prophage group but different integrase type and *vice versa*.

To provide a broader context to the prophages from Argentinian GBS, a phylogenetic analysis of 764 prophages from GBS (isolated in Argentina and worldwide) and other 43 streptococcal species was performed (Table S3 in S2 File).

### M6. Integration of information

The Microreact application [27] was used for an integral visualisation of the collected information.

### M7. Statistical analysis

Fisher’s exact test (two-tailed) was used to evaluate the association between prophage presence and GBS clonal complex. A *p*-value of ≤0.05 was considered to be significant.

### M8. Ethics statement

Ethical approval for the ‘‘Argentinian multicentric study on infections due to *Streptococcus agalactiae*’’ was provided by the Ethics Committee of the Faculty of Pharmacy and Biochemistry, University of Buenos Aires, Res [D] RESCD-2022-400-E-UBA-DCT.

## Results

### R1. Prophage detection and typing in GBS genomes from Argentina

Prophage screening based on prophage phylogenetic group and integrase type, [11,16] detected 383 putative prophages in the 365 GBS genomes. A total of 200/383 (52%) prophages were grouped into 10 prophage types (prophage group + integrase type). In 60/383 (16%) prophages only the integrase type was determined (the prophage group was not determined [ND]). A total of 123/383 prophages (32%) belonged to prophage group A, known to lack the integrase gene [16,28].

In contrast, a manual search revealed 325 prophages among the 365 GBS genomes. Forty-two out of the 325 prophages (13%) were fragmented in two or more contigs but were successfully assembled with reference-based read mapping. The additional prophage sequences detected based on the screening approach (n=58) represented mostly groups A (36/58 lacked the lysis module) and F (19/58 integrase gene only present). In 3/58 cases the screened prophages were disregarded due to high level of fragmentation.

Only the 325 manually detected prophage sequences were analysed further. Phylogenetic analysis revealed clustering by the prophage group (Fig 3, Fig S1A in S3 File). Prophages that could not be typed by group with the initial screening (ND prophages) were assigned a prophage group corresponding to their phylogenetic cluster. Prophage groups E and F were further divided in two subclusters each (100% bootstrap support). In both cases, prophages of one subcluster had not been detected by the *in silico* PCR. Those that were detected by the *in silico* PCR were reclassified as subgroups E1 and F1, respectively, while those not detected were reclassified as subgroups E2 and F2, respectively (Fig S1B in S3 File). The reclassification of group F prophages into the subgroups F1 and F2 coincides with van der Mee-Marquet’s classification of prophages with insertion sites F1 and F2, respectively [11].

**Fig 3.**
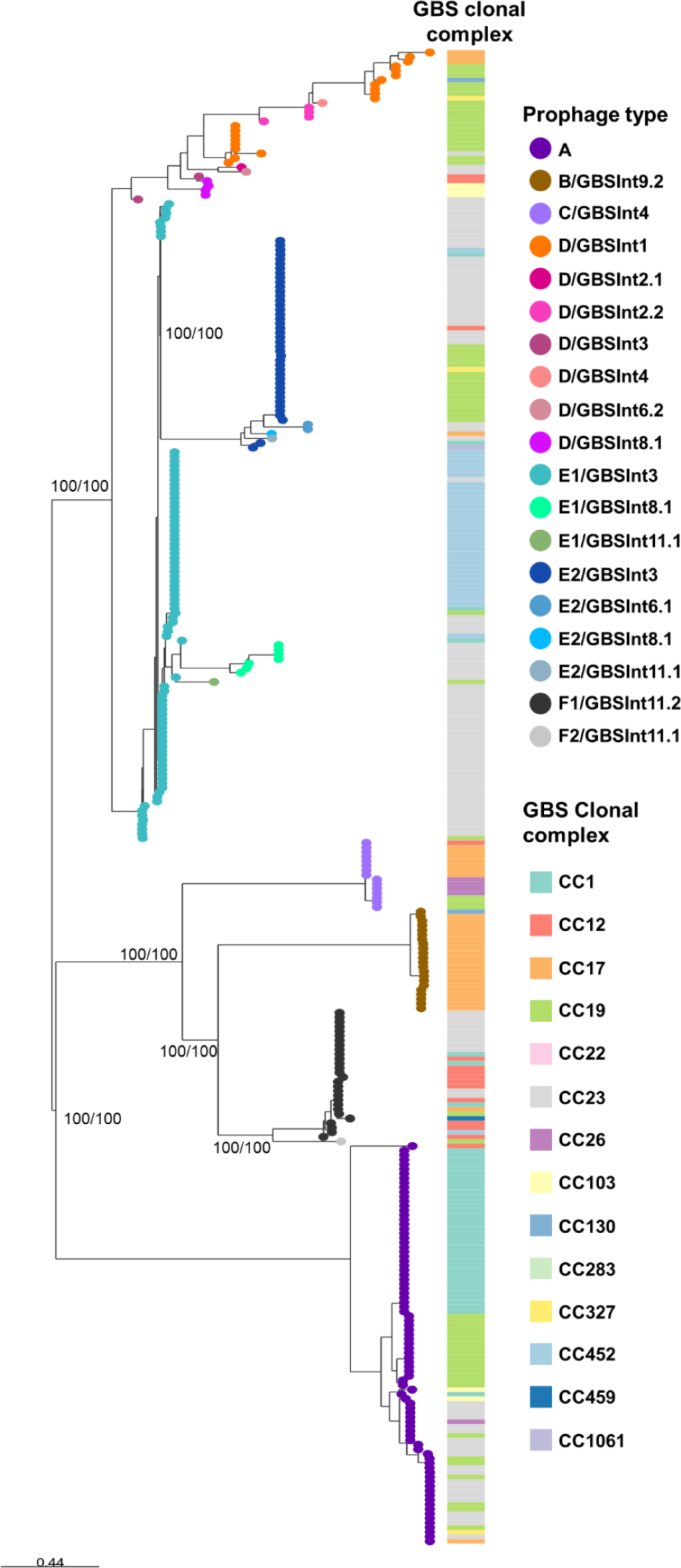
Phylogeny of 325 prophages found in 365 Argentinian Group B *Streptococcus* (GBS) genomes. Maximum-likelihood phylogenetic tree, midpoint rooted, with nodes coloured by prophage type (determined based on the combination of prophage group and integrase type). Support values (SH-aLRT/ Ultrafast Bootstrap) are shown as labels for selected nodes. Host GBS clonal complex is shown as coloured blocks. The scale bar represents the number of SNPs per variable site. https://microreact.org/project/philogeny-argentinean-gbs-prophages

As a result of this integrated analysis all prophages (n=325) were reclassified into 19 prophage types (Fig 3). The majority of GBS genomes (54%) carried a single prophage. Multiple integrase types/subtypes were found within prophage groups and vice versa (Fig 3), as previously described [16], indicating that screening of GBS prophages by previous methods alone is insufficient to accurately classify the prophages.

### R2. Improvement of the screening and typing system

In order to improve the screening and typing system for GBS prophages, we developed a database of phylogroup-specific prophage genes (Table S1 in S2 File, Data S1 in S3 File). The genes were selected based on 12 PCR-amplified fragments using previously described primers [11], and refined to avoid false positive results. Detection of group A prophages is based on genes *hha*I or *clp*P combined with a presence of either a holin or a lysin gene (Table S1 in S2 File). F1 prophage integrase gene (*hin*) was replaced by a gene coding for a terminase large subunit (Table S1 in S2 File). The hypothetical prophage gene representative of group D prophages was replaced by two new genes (Table S1 in S2 File). Finally, gene sequences for the detection of group E2 and F2 prophages were also added to the database (Table S1 in S2 File). These changes markedly improved the specificity and accuracy of prophage detection and typing in the 365 GBS genomes from Argentina (Fig 4).

**Fig 4.**
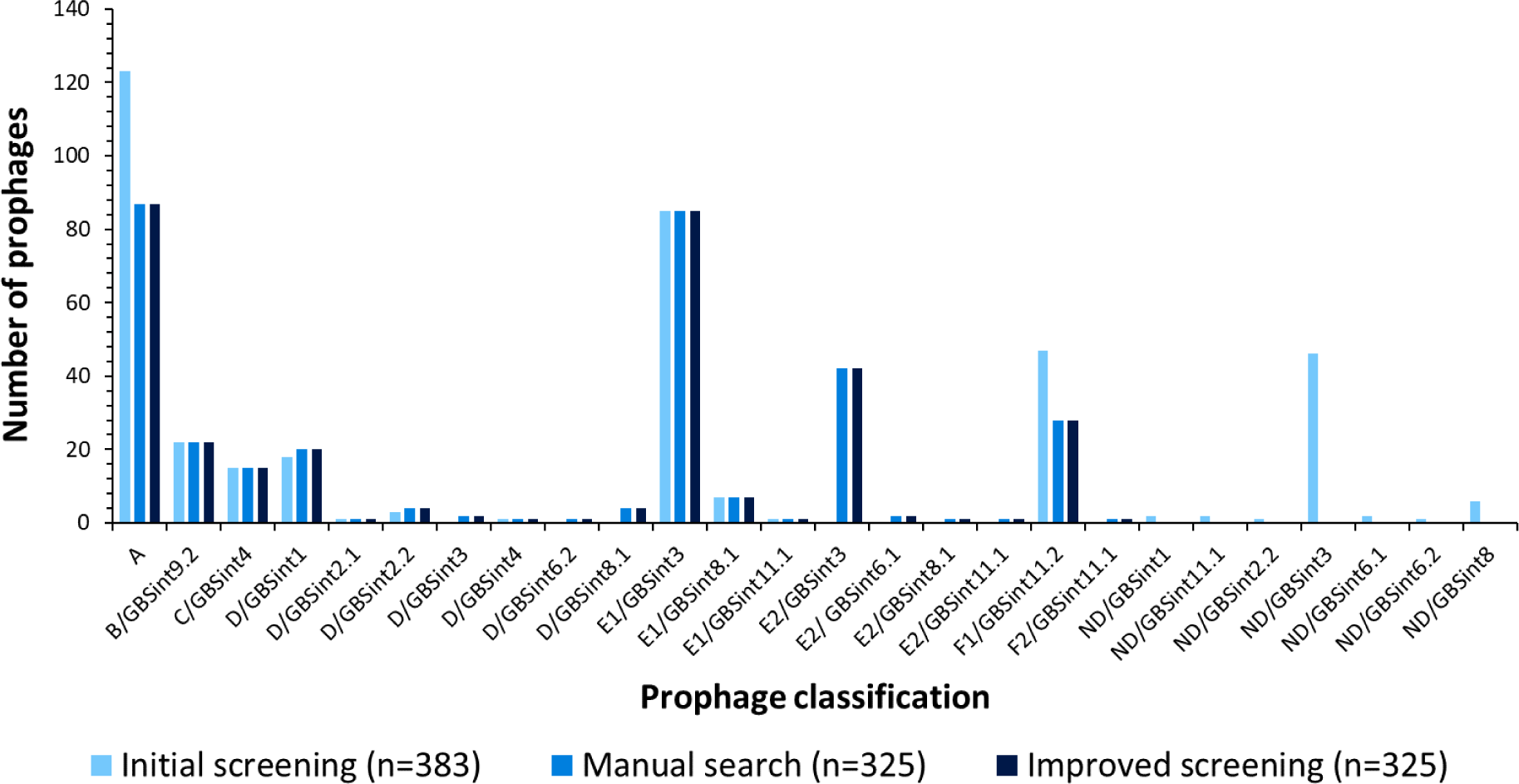
Prophage detection and typing in 365 Argentinian Group B Streptococcus (GBS) genomes. Three methods are compared: the initial integrated screening of prophage group by *in silico* PCR (with primers designed by van der Mee-Marquet et al.) and integrase type by blastx search against a GBS-prophage integrase database (built by Crestani et al.); the manual search in the GBS genomes of the screened prophages and their classification according to their phylogeny; the improved screening and typing method by prophage group: blastn search against a database of the representative genes for each group proposed by van der Mee-Marquet et al., and new genes proposed in this work, integrated with the integrase typing designed by Crestani et al.

More importantly, our improved method detected 10 additional prophage types when tested on a collection of 615 globally-derived GBS genomes (including the 365 from Argentina), giving a total of 29 distinct prophage types. The 10 prophage types not found among the GBS genomes from Argentina included 2 new integrase subtypes: GBS*Int*6.3 and GBS*Int*8.2 and their sequences were added to the integrase database (Data S2 in S3 File).

A graphical summary of the proposed improved screening and typing method for GBS prophages is shown in Fig 5.

**Fig 5.**
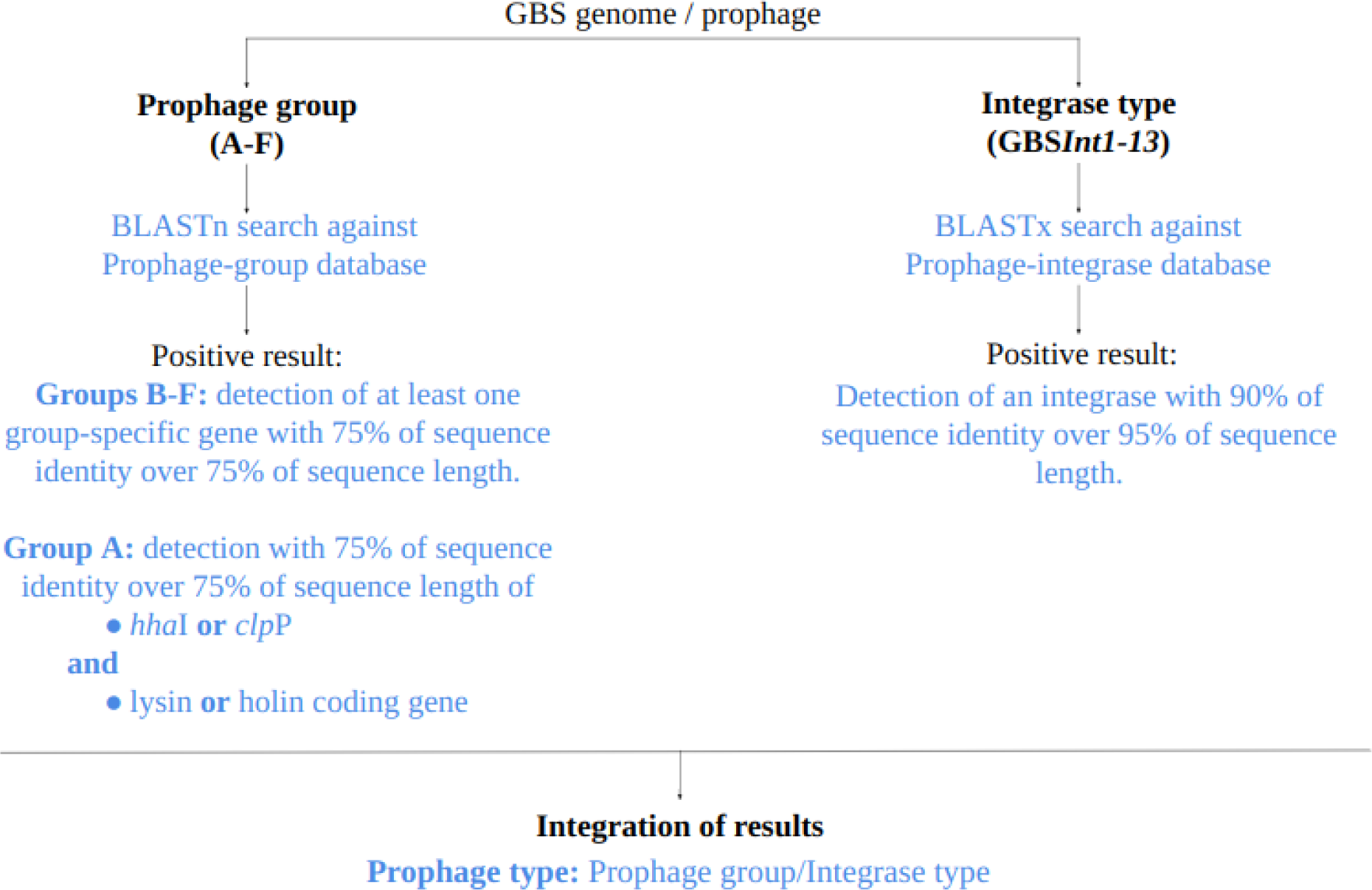
Methodology summary for the improved screening and typing system for GBS prophages.

### R3. Characterization of GBS prophages from Argentina

The most prevalent prophage types in GBS isolates from Argentina were: A (87/325, 27%), E1/GBS*Int*3 (85/325, 26%), E2/GBS*Int*3 (42/325, 13%) and F1/GBS*int*11.2 (28/325, 9%), (Fig 4). Significant associations were found between most prophage types and CC assignment (*p*<0.05, Fig S2 in S3 File). Prophages of type A were associated with CC23 and CC1; B/GBS*Int*9.2 with CC17; C/GBS*Int*4 with CC17 and CC26; D/GBS*Int*1, D/GBS*Int*2.2 and E2/GBS*Int*3 with CC19; D/GBS*Int*8.1 with CC103; E1/GBS*Int*3 with CC23 and CC452; E1/GBS*Int*8.1 with CC23; F1/GBS*Int*11.2 with CC12.

The phylogenetic analysis (Fig 3) revealed monophyletic clusters for most prophages within a group or subgroup with the exception of group D prophages, which were more diverse.

The insertion site and *att* sequences matched those described by Crestani et al. [16] for each integrase type, with the exception of some prophages with minor changes in their *att* sequences (Table S4 in S2 File). One gene of each prophage module (integrase, helicase, terminase large subunit, major capsid protein, lysin) was selected for phylogenetic analysis of their nucleotide sequences, to determine whether similar modules could be found in different prophage types. These analyses showed, in general, similar clustering between phylogenies based on individual prophage module genes and that observed based on alignment of full prophage sequences (Fig S3 in S3 File).

Several genes potentially beneficial for the host bacteria were found exclusively in B/GBS*Int*9.2 prophages, including genes involved in carbohydrate or RNA metabolism, which were present in all prophages of this type. A single B/GBS*Int*9.2 prophage additionally carried genes coding for phosphoenolpyruvate synthase, a multidrug resistance efflux pump, genes involved in DNA metabolism and several genes encoding different types of permeases (Table S5 in S2 File).

Phylogenetic analysis of 764 *Streptococcus* spp. prophages revealed that GBS phages from Argentina were related to other GBS prophages and with bacteriophages from other streptococcal species (Fig 6). In particular, group B prophages were closely related to *S. pyogenes, S. iniae, S. oralis* and *S. pneumoniae* phages, while group A and F prophages were related to phages from more than ten different streptococcal species. However, prophages from groups C, D, and E were more closely related to each other in this phylogeny (100% bootstrap support) and seem to be mainly restricted to GBS, with an occasional spread to *S. dysgalactiae, S. equi, S. iniae* and *S. pyogenes* (clustering with group C) and one single *S. uberis* prophage (clustering with group E1).

**Fig 6.**
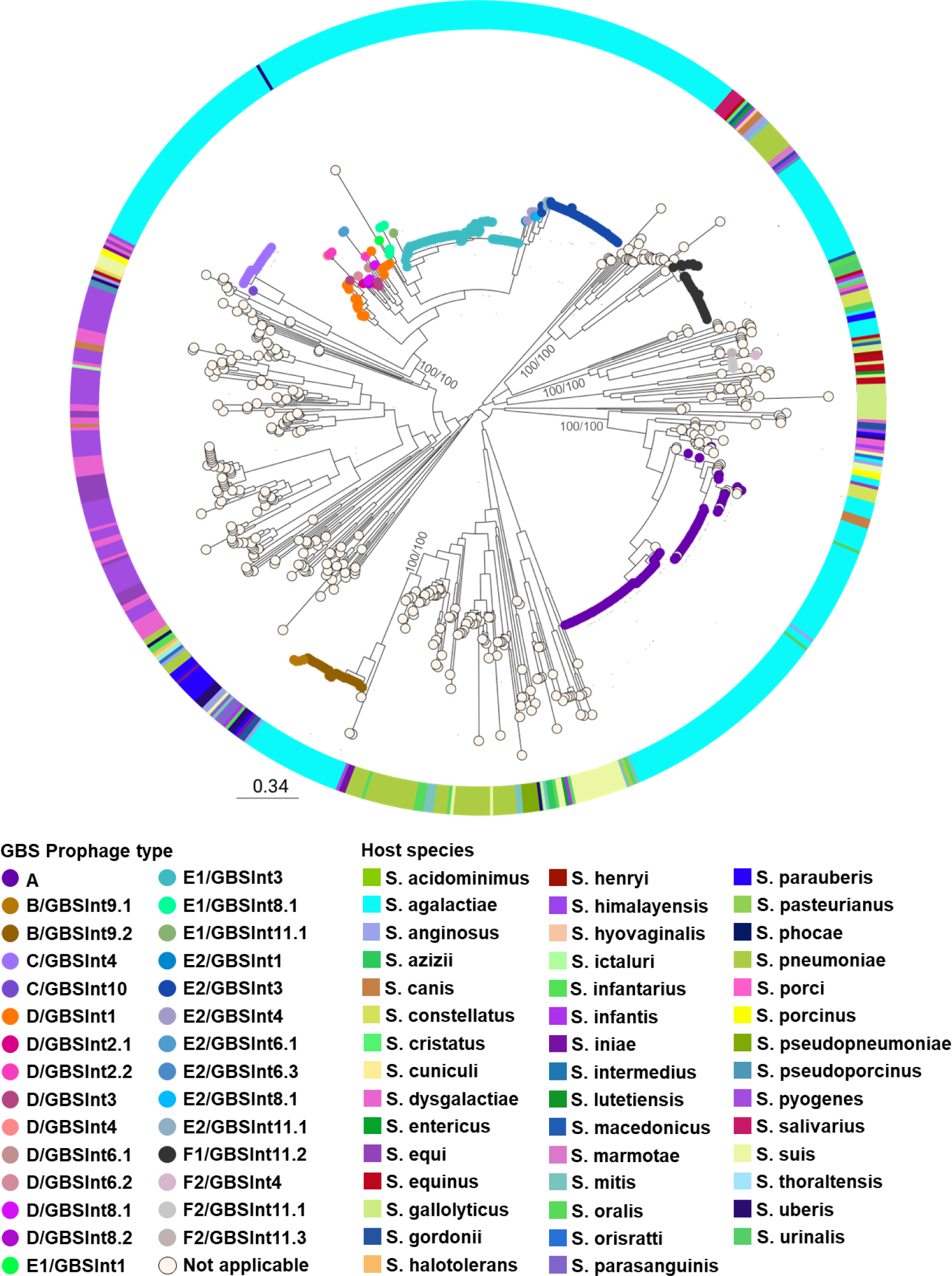
Phylogenetic tree of prophages from streptococcal species. Maximum-likelihood phylogenetic tree of 764 prophages found in 580 genomes from 44 streptococcal species, including GBS. Nodes are coloured by GBS prophage type, where applicable. Host species are shown in the coloured ring. Tree was rooted at midpoint. Support values (SH-aLRT/ Ultrafast Bootstrap) are shown as labels for selected nodes. The scale bar represents the number of SNPs per variable site. https://microreact.org/project/gbs-prophages-in-a-global-context

### R4. Comparative analysis of the 29 distinct prophage types

More than one integrase type or subtype was found in all prophage groups, with the exception of group A (prophages without integrase). Integrase types GBS*Int*1, GBS*Int*3, GBS*Int*4, GBS*Int*8.1 and GBS*Int*11.1 were found in more than one prophage group, mostly D and E. Also, D and E2 prophages had the most diversity in integrase types (7 and 6 types, respectively). Group B and F prophages had two subtypes of the same integrase type (GBS*Int*9 and GBS*Int*11, respectively).

One prophage sequence for each of the 29 types was selected for further characterization. Based on their predicted morphology, all phages were classified as siphovirus (former *Siphoviridae* family). Visual inspection of the annotated prophage sequences confirmed the modular organisation following a specific order based on their function in the phage life cycle: lysogeny, replication, packaging, morphogenesis and host lysis (Fig 7). Genome sizes ranged from 32.6 to 47.9 Kb, with group B prophages the smallest (likely due to shorter genes and a shorter morphogenesis module) and the prophages of groups D and E1 the largest (likely due to higher gene content in the lysogeny and replication modules). Manual inspection of the sequences revealed that all prophage types contained open reading frames (ORFs) encoding putative proteins related to: bacterial fitness, defence mechanisms and/or virulence (Table S4 <u>in S2 File</u>). ORFs coding for hypothetical proteins where no known conserved domain was found constituted from 29 to 59% of the coding sequences of each prophage. These genes were found throughout the entire prophage genome and not organised in a single module, although they were more frequent in the lysogeny and replication modules.

**Fig 7.**
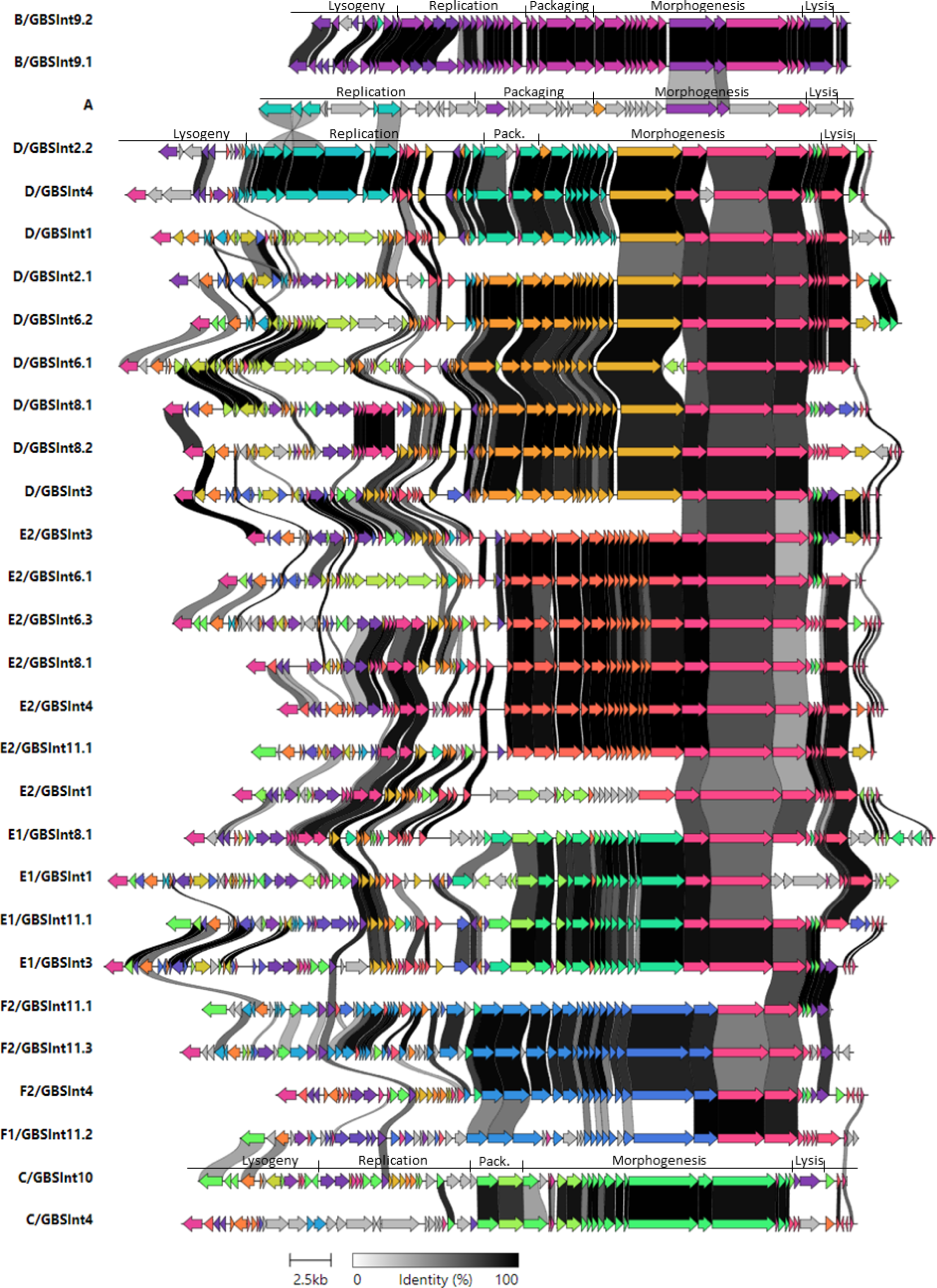
Comparative analysis of representative sequences of 29 distinct prophage types found in the analysed GBS. Colours represent groups of homologous genes. Genes with more than 40% of identity are linked with grey-black strokes, as shown in the scale.

The amino acid sequences of each integrase type and lysins found in each of the 29 reference prophages, and lysins coded by genes from each phylogenetic cluster (Fig S3E in S3 File) were analysed to determine their catalytic domains (Table S4 in S2 File). All integrase types had a tyrosine recombinase domain, except for GBS*Int*11.1 and GBS*Int*11.2, which had a serine recombinase domain. The putative lysins were classified based on their cleavage site, into three of the five major endolysin classes [29]: N-acetyl-β-D-glucosaminidase, N-acetyl-β-D-muramidase and N-acetylmuramoyl-L-alanine-amidase. Interestingly, lysins from all prophages contained the domain N-acetylmuramoyl-L-alanine-amidase and, in most cases, the lysins were bifunctional, as they also carried a second catalytic domain with a different cleavage site, either N-acetyl-β-D-glucosaminidase or N-acetyl-β-D-muramidase. All prophages from the same type had lysins of the same class, except for types A, C/GBS*Int*4, D/GBS*Int*3, E1/GBS*Int*3, E2/GBS*Int*4, E2/GBS*Int*11.1 and F1/GBS*Int*11.2, in which some lysins had only the N-acetylmuramoyl-L-alanine-amidase domain, while others were bifunctional.

Comparative sequence analysis of the 29 prophage types (Fig 7) showed more than 60% identity between the morphogenesis modules encoding most tail proteins of the groups D, E and F and around 45% identity between most genes in the replication modules of the type A, D/GBS*Int*4 and D/GBS*Int*2.2. Group F prophages shared more similarities in a modular level with groups D and E than with phylogenetically closer groups B and C, which did not show identity on a modular level with other prophage types.

Comparative analysis of prophages belonging to the same prophage group but carrying different integrase types or subtypes (Fig S4 in S3 File) revealed that the major differences between the prophages within groups B, C, D and E and subgroup F2 resided in the lysogeny and replication modules. The F1prophage showed less than 60% identity with subgroup F2 between all their modules except for the morphogenesis. In addition, prophages with the same integrase type but belonging to different prophage groups (Fig S5 in S3 File) did not reveal any similarity, other than the integrase gene itself. The exceptions were prophages with integrase type GBS*Int*4, where more than 90% of identity was found between the lysogeny module and the genes following the host lysis module (Fig S5C in S3 File).

## Discussion

Prophage presence has been reported to impact GBS epidemiology in collections from Europe [11,13–15]. In particular, the acquisition of certain prophages has been linked with the emergence of GBS infections in neonates and adults in France [11] and it has been postulated that the presence of prophages encoding virulence factors was responsible for the increased incidence of severe neonatal infections both in France [13,28] and the Netherlands [14].

This is the first report on the diversity of prophages in human GBS isolates from South America. Employing two methods for GBS prophage typing [11,16] and subsequent manual inspection, we were able to detect 325 prophages among 365 GBS isolates collected in an Argentinian multicentric study.

### New prophage typing system

We propose an enhanced method that combines and simplifies some screening steps of previously developed GBS-prophage typing methods that allowed the screening and classification of prophages by either phylogenetic group [11] or integrase type [16], which do not often correlate [16, this work]. Our results show that using each method individually can lead to false negative or false positive results. Furthermore, it can be insufficient to classify GBS prophages accurately, as determining the prophage group provides information about the genomic characteristics of the phage but not its insertion site. The latter can be determined based on integrase type, which in contrast does not offer insights into composition of full prophage sequence.

The new method increases detection of full prophage sequences, as well as prophages that are fragmented into multiple contigs. Additionally, it allows the classification of prophages based on both their phylogenetic lineage and integration site within the bacterial genome. Using this approach, we were able to identify a total of 29 distinct prophage types, including 19 prophage types in GBS from Argentina and 10 additional types in prophages from GBS collected in other countries.

Our results demonstrate that this improved integrated method is less likely to detect prophage remnants, allows identification of novel prophage integrases and offers fast detection of GBS prophages in a large genomic dataset, with little computational processing.

### Evolution of GBS prophages

Genome mosaicism was observed in all prophage types, in accordance with the proposed modular evolution of prophages [30–32]. Genes belonging to the packaging module, those encoding capsid proteins and a few coding for tail proteins appear to have been acquired as a block, independently of the rest of the prophage genes. This is especially evident in groups with a greater number of prophage types, D and E, since the clustering of phages into phylogenetic subclades (Fig 3) correlates with their grouping according to the homology in the aforementioned gene region (Fig 7). The presence of homology between several genes encoding tail proteins in prophages D, E and F suggests that this group of genes share a common ancestor and that they might have been acquired in an independent recombination event from the rest of the structural genes, which did not present homology between the different phage groups (Fig 7). In general, prophages within the same group presented the same classes of lysins (same catalytic domains, Table S4 in S2 File), even if they did not share high homology in their lysis modules. However, there was no lysin class exclusive to one prophage group, which could indicate an independent recombination of the lysin-coding gene from the rest of the lysis module.

Divergence between prophages, even belonging to the same phylogenetic group, was typically observed in the lysogeny and replication modules, where the majority of the genes encoding hypothetical proteins and genes potentially beneficial for GBS were located. This suggests that these regions would be the most prone to suffer recombination events. This is also supported by the lack of similarities in the lysogeny module of prophages with the same integrase type but different prophage group, which implies that the prophage integration site in the host genome is dependent only on the integrase gene and *att* sequence. The general absence of homology between prophages of groups A, B and C with other prophage types would suggest that these prophage groups have little propensity for horizontal gene exchange with GBS prophages from other groups.

Most prophage groups and subgroups seem to be highly clonal, whereas group D showed more divergence, demonstrating evidence of microevolution (Fig 3, Fig 6 and Fig S1A in S3 File). The reason for such variability is not clear yet, but future studies are needed to understand if it might be advantageous to the host.

The level of conservation of genes encoding tail proteins among various prophage groups (Fig 7) could mean that they are involved in the specific recognition between the phage and the bacterial receptor, a process that has not yet been studied in GBS. If so, these phages could have a similar host range. Interestingly, tail protein genes are also conserved in prophage types from other streptococcal species (*S. pyogenes*, *S. pneumoniae*) [33,34]. Further analysis of the tail protein sequence conservation can reveal new insights into the mechanisms of prophage sharing between *Streptococcus* spp.

### Prophage presence in the context of GBS epidemiology

The prevalence of GBS isolates carrying at least one prophage and the average number of prophages per isolate found in our dataset correlates with previous reports on prophage content in GBS [11,13,15,35,36]. Our results also aligned with the integrase type and CC associations reported previously by Crestani [16]. The reason for such association should be explored, but could be related to the presence of specific restriction-modification systems, which may limit the recombination and horizontal transfer of mobile genetic elements between GBS lineages, as seen in other bacterial species [37–39].

In line with previous reports, GBS prophages identified in our dataset carried genes potentially involved in bacterial fitness, host adaptation to stressful environments and virulence [11,13,15,28,40]. However, most of these genes were not detected by the online prophage genome screening tools, which stresses how manual curation remains an important part of the prophage genome investigations. A single B/GBS*Int*9.2 prophage was found to carry a gene encoding a multidrug and toxic compound extrusion (MATE)-like protein, similar to MepA from *Staphylococcus aureus* [41]. It has been reported that expression of MATE proteins in bacteria confers resistance to toxic dyes and multiple antibiotics [42,43].

While carriage of antibiotic resistance genes (ARGs) is not common in prophages [44], there have been reports of ARGs carriage in conjugative elements within prophages of streptococcal species [45,46]. The fact that these elements were found in only a single GBS prophage analysed in this study, highlights the overall low prevalence of ARGs in GBS prophages. However, it is important to note that, on average, 50% of GBS prophage genes code for hypothetical proteins lacking recognizable conserved domains. Hence, it is important to continue the study of these prophages to better understand their biology and impact on the host cells, particularly in a context of bacteriophage therapy development [47].

GBS carriage of prophages associated to those from other streptococcal species causing infections in humans and animals has been extensively documented in the literature [11,15,35,48,49]. Our results further show that prophages detected in GBS isolates from Argentina are globally distributed and suggest that prophages belonging to groups A, B, and F might have evolved from prophages horizontally transferred between different species of streptococci (Fig 6). In contrast, prophage groups C, D and E, that were found to share a common ancestor in this global phylogeny, seem to be mainly restricted to GBS (Fig 6).

Further experimental work is needed in order to confirm the prophage transfer between streptococcal species, receptor specificity (tail proteins) and the restriction of C, D, and E prophages to GBS only. It is also crucial to investigate if the insertion sites of these prophages favour the mobilisation of genetic material and the potential for horizontal transfer of genes present in defective prophages or phage remnants.

### Study limitations

Limitations of this study include using GBS genome assemblies from short read data to detect and classify prophages. All prophage sequences identified after manual curation remain theoretical, even when found in the same contig, due to possibility of assembly errors, particularly in prophage modules with high sequence similarity between different phage types (e.g., structural modules). This could be addressed by performing long-read sequencing followed by a hybrid assembly.

### Conclusions

This study performed a comprehensive comparative analysis of prophage genomes in GBS isolates from Argentina, and is a first report on GBS prophage diversity in Latin America. We propose the use of an improved and integrated prophage typing system suitable for rapid phage detection in GBS genomes and their classification with little computational processing. The presence of prophages in most GBS isolates analysed here, association between prophage groups and GBS lineages as well as carriage of genes beneficial for the host bacteria reinforce the hypothesis that acquisition of prophages confers an evolutionary advantage to GBS and may play an important role in its epidemiology. The diversity of prophage types found in GBS isolates from Argentina along with the observed lysin diversity is a promising finding, which can be explored further to identify novel lysins with activity against GBS as an alternative therapy for GBS infections. In the light of the global challenge posed by antimicrobial resistant bacteria it is imperative to advance our current knowledge of bacteriophage biology and applicability of lysins as an alternative treatment against bacterial infections, to promote and expedite the approval and regulation of such therapies.

## Acknowledgments

We would like to thank everyone involved in the genome sequencing of our GBS collection at the sequence facilities of the Wellcome Sanger Institute and Instituto Malbrán.

## Supporting information

**S1 File. Detailed methodology**

**S2 File. Supplementary tables**

**S3 File. Supplementary figures and data**

